# Proton Transfers to DNA in Native Electrospray Ionization Mass Spectrometry: A QM/MM Study

**DOI:** 10.1101/2022.10.10.511116

**Authors:** Mirko Paulikat, Juan Aranda, Emiliano Ippoliti, Modesto Orozco, Paolo Carloni

**Author notes:** **Corresponding Authors** Paolo Carloni – *Computational Biomedicine (IAS-5/INM-9), Forschungszentrum Jülich GmbH*, *52428 Jülich, Germany*; *Department of Physics, RWTH Aachen University, Otto-Blumenthal-Straße, 52062 Aachen, Germany*;, Modesto Orozco – *Institute for Research in Biomedicine (IRB) Barcelona, The Barcelona Institute of Science and Technology, 08028 Barcelona, Spain*; *Department of Biochemistry and Biomedicine, University of Barcelona, 08028, Spain.

## Abstract

Native electrospray ionization - ion mobility mass spectrometry (N-ESI/IM-MS) is a powerful approach for low-resolution structural studies of DNAs in the free state and in complex with ligands. Solvent vaporization is coupled with proton transfers from ammonium ions to the DNA resulting in a reduction of the DNA charge. Here we provide insight on these processes by classical MD and QM/MM free energy calculations on the (GpCpGpApApGpC) heptamer, for which a wealth of experiments is available. Our multiscale simulations, consistent with experimental data, reveal a highly complex scenario: the proton either sits on one of the molecules or is fully delocalized on both, depending on the level of hydration of the analytes and on size of the droplets formed during the electrospray experiments. This work complements our previous study on the *intramolecular* proton transfer on the same heptamer occurring after the processes studied here, and, together, provide a first molecular view of proton transfer in N-ESI/IM-MS.

**TOC GRAPHICS:** 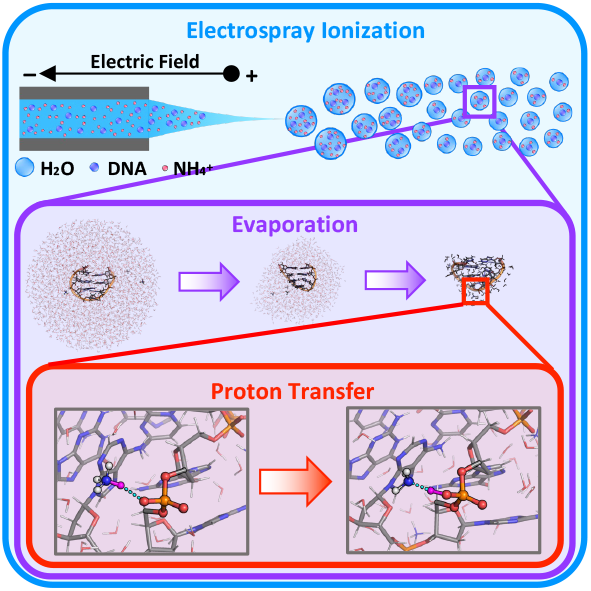

---

Native electrospray ionization - ion mobility mass spectrometry (N-ESI/IM-MS) is a very powerful tool for low-resolution structural studies of DNA oligomers and their complexes with ligands.^1–4^ It requires much smaller quantities than high resolution techniques, such as NMR, X-ray crystallography or cryo-electron microscopy.^5,6^ The biomolecular analytes are sprayed from solution in quasi-physiological conditions (aqueous ammonium solutions at neutral pH, without organic cosolvents)^7,8^ into the gas phase through a capillary applied to an electric field.^9,10^ Water droplets are formed and eventually evaporate up to a complete loss of solvation.^11^ Originally, ammonia is positively and phosphates are negatively charged, but as solvent evaporates the ammonium ions transfer protons to the DNA phosphates, consistently with the gas phase basicities of NH_3_ (819 kJ· mol^−1^)^12^ and dimethyl phosphate (DMP; 1,360 kJ· mol^−1^).^13^ This leads to a decrease in absolute value of the analyte’s charge (along with a loss of NH_3_), in turn associated with a high signal-to-noise ratio.^14^

Molecular simulation (MD) studies have provided important insight on DNA oligomers in the gas phase.^15^ They suggested that duplex,^16^ triplex,^17^ G-quadruplex^18^ oligomers maintain several structural determinants on passing from the solution to the gas phase. They have also described the impact of the vaporization process in the structural ensemble.^19^ Furthermore, when combined with quantum calculations, MD simulations show that once protons are transferred to the DNA, they can jump from different basic sites in the oligos, leading to some structural changes and to a complex fuzzy change density in the analyte.^20^ The remaining question is then how the intermolecular proton transfer events, from ammonium to DNA, occur during vaporization and what is their structural impact.

Here we perform molecular simulations on d(GpCpGpApApGpC) heptamer and ammonium ions in water and gas phase to investigate ammonium-DNA proton transfers. First, we simulate the evaporation process under native ESI-MS conditions by batches of classical MD simulations of the heptamer immersed in a water droplet containing ammonium ions. We remove dissociating water molecules progressively. Second, we study the proton transfer phenomena between the ammonium ion and the heptamer -represented either as DMP (**I**, Fig. 2) or in its integrity (**II**, Fig. 4)- in the partially dehydrated droplets. To this aim, we use quantum mechanics/molecular mechanics (QM/MM) MD and umbrella-sampling-based free energy simulations, exploiting QM/MM interfaces recently developed with the support of the EU projects BioExcel and BioExcel-2 (www.bioexcel.eu).

## MD simulations of complex I

Here, the heptamer is represented as dimethyl phosphate (DMP). First, we run 10ns-long force field-based simulations at 300 K and 1 bar of ammonium and DMP (as ionic moieties) in water. Both are fully solvated, as shown by the plots of radial distribution functions (rdf, Fig. S1). Integration of the latter shows that the average number of waters surrounding the ammonium ion (hydration number, *HN*) is 8, and the DMP solvation structure shows 4–5 water molecules in the first shell for each anionic oxygen atom. The total hydration number of DMP, estimated from the oxygen-phosphate rdf, counts 10 water molecules (Fig. S1). The distance between ammonium nitrogen and DMP phosphorus atom (*d*) varies widely from 4 (contact ion pairs) to 30 Å (size of the simulation box). The ion pairs are of very transient nature, lasting typically less than 1 ns (Fig. S2).

Next, we let three droplets containing ∼800 water molecules (extracted from selected MD snapshots) evaporate by MD simulations at 300 K. Although most waters dissociate quickly from the droplet (Fig. 1a, Fig. S3), the ions remain fully solvated until ∼45 ns (8 coordinating water molecules). Further evaporation leads to only 3 coordinating water molecules (Fig. 1b). The last water molecule dissociates after 102 ns at an increased temperature of 450 K. The distance *d* decreases as the droplet shrinks; transient ion pairs are formed several times during the dynamics. After ∼45 ns, the two ions form direct H-bonds (Fig. 1c).

**Figure 1:**
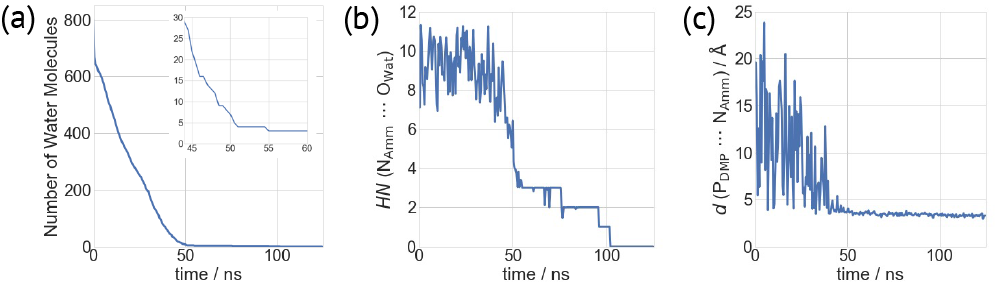
Gas phase MD simulations of complex **I**: different quantities plotted as a function of simulated time: (a) Number of water molecules present in the droplet. Inset: Enlarged view from 45 ns to 60 ns, when permanent ion pairs are formed. (b) Hydration number (*HN*) of the ammonium ion. (c) Distance *d* between the DMP-phosphorus atom (P_DMP_) and the NH_4_^+^-nitrogen atom (N_Amm_).

### QM/MM free energy calculations of complex I

After 2.4 ps-long QM/MM MD re-equilibration of the system (see SI, Fig. S4), we carry out US free energy calculations at 300 K with different water content, from 3 (**Ia)** to 4 (**Ib)** and 8 **(Ic)** (Fig. 2). The collective variable (CV) used here is the difference between the N–H breaking bond distance and the O–H forming bond distance (Chart S1): negative and positive values of the CV are associated with ionic and neutral states, respectively. The free energy profile is well converged, as shown by its time evolution (Fig. S5).

**Figure 2:**
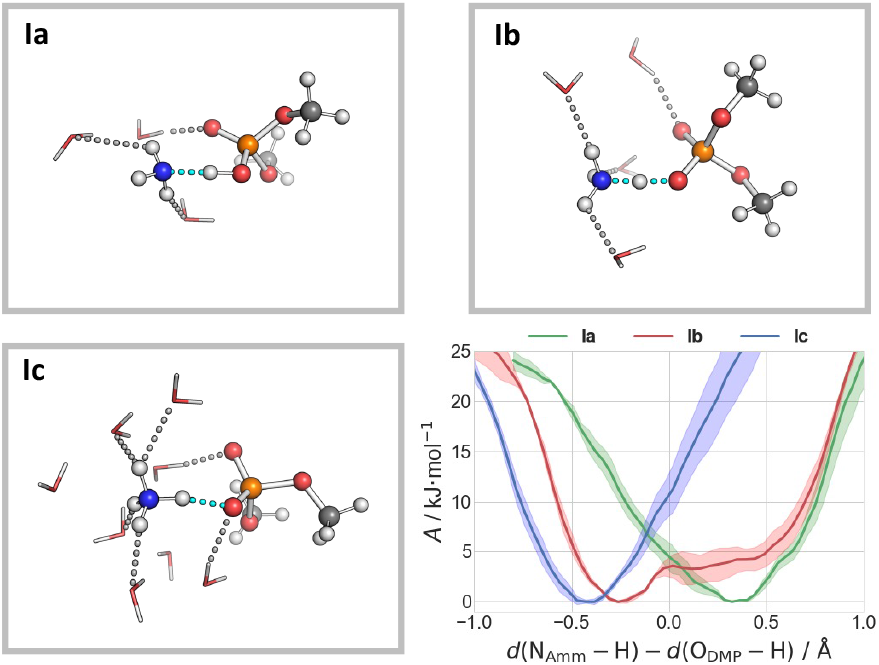
Snapshots of the three systems **Ia, Ib**, and **Ic**, indicating the preferred protonation state of the ammonium and DMP moieties. H-bonds are shown as dashed lines. Lower right: Free energies as a function of the difference of the breaking/forming N_Amm_–H and H–O_DMP_ bond distances.

In **Ia**, the proton is localized on DMP with a poor hydration of the ammonia moiety (*HN* of ∼2– 3, see Fig. 2). By slightly increasing the hydration *(HN* ∼ 3.5) the proton is shared by the two moieties (**Ib**). Further increasing the hydration (*HN* ∼ 5 in **Ic**) leads to the stabilization of the ammonium ion (Fig. 2). Thus, the protonation state is very sensitive to the local solvation environment: only if the ammonium is largely solvated does the cation keep the proton.

### MD simulations of complex II

A 500ns-long classical MD simulation, in the same conditions as above, is carried out on d(GpCpGpApApGpC) and six ammonium ions in aqueous solution. The overall charge of the system is zero.

The results are similar to previous studies on the biomolecule alone in water (see SI).^21,22^ In particular, the oligonucleotide retains its compact hairpin structure, which consists of a short B-DNA segment and a sharp turn within the d(G_3_A_4_A_5_)-triloop with one non-canonical d(G· A) base pair.^21,22^ The radius of gyration (*R*_gyr_) is 7.05 ± 0.06 Å (Fig. S6–7). Three and two H-bonds are observed regularly for the two canonical base pairs (G_1_–C_7_ and C_2_–G_6_) and the non-canonical base pair G_3_–A_5_, respectively.

The two ionic moieties are well separated: Only 1.4% of these are transient ion pairs (Fig. S6), with an average hydration number equal to 8 for both species (Fig. S1).

Three water droplets with three ammonium ions (Fig. S8, total charge of –3*e*, the main charge state observed in ESI-MS experiments),^20^ extracted by the MD simulations, underwent 225 ns of classical MD simulations. As all simulations showed very similar features, only one is discussed here (details for the others in the SI, Fig. S9–S13). During the first 150 ns (at 300 K), several water molecules strip out (Fig. 3a), but the oligomer maintains its intramolecular H-bonds as in water solution (Fig. S11–S13). The ions are fully solvated within the first 100 ns, but in the next 50 ns, the ammonium *HN* decreases from 8 to 2. Simultaneously, the oligomer becomes more compact (*R*_gyr_ ∼ 6.9 Å compared to ∼ 7.05 Å in solution). The two ions get closer as the water droplets shrink, as observed for **I** (Fig. 3d). The ammonium ion forms two types of direct H-bonds with the phosphate moieties, associated with different hydration numbers (Fig. 3e): with only one phosphate group (interaction mode **(P)**, Fig. 4a-b and Fig. S14a) or with two phosphate groups from adjacent nucleotides (interaction mode **(P2)**, Fig. 4c-d and Fig. S14b). After ∼150 ns, about 30 water molecules are present. By increasing the temperature up to 450 K, the last water molecules evaporate. The oligomer becomes even more compact (*R*_gyr_ ∼ 6.8 Å). The canonical and non-canonical base pairs may dissociate forming other hydrogen bonds (see Fig. S11–S13). Such compact conformation is similar to that of the same system without the ammonium group.^20^

**Figure 3:**
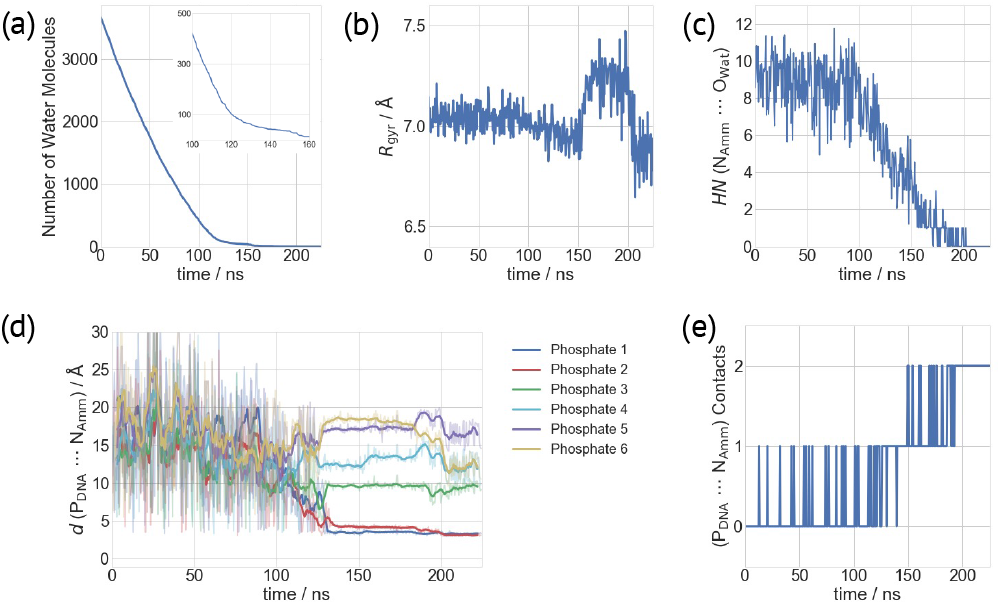
Gas phase MD simulations of complex **II** – different quantities plotted as a function of simulated time: (a) Number of water molecules. Inset: Enlarged view from 100 ns to 160 ns. (b) Radius of gyration (*R*_gyr_) of the DNA oligo. (c) *HN* values of NH_4_^+^-nitrogen atoms (N_Amm_) surrounded by water-oxygen atoms (O_Wat_). (d) Distances of the DNA-phosphorus atoms (P_DNA_) and one N_Amm_-atom. Data at the end of each 500ps-long batch of MD simulations are shown as transparent lines, while their moving averages (window size of 10) are represented as bold lines. (e) Number of contacts of one NH_4_^+^ ion with the DNA phosphate groups.

**Figure 4.**
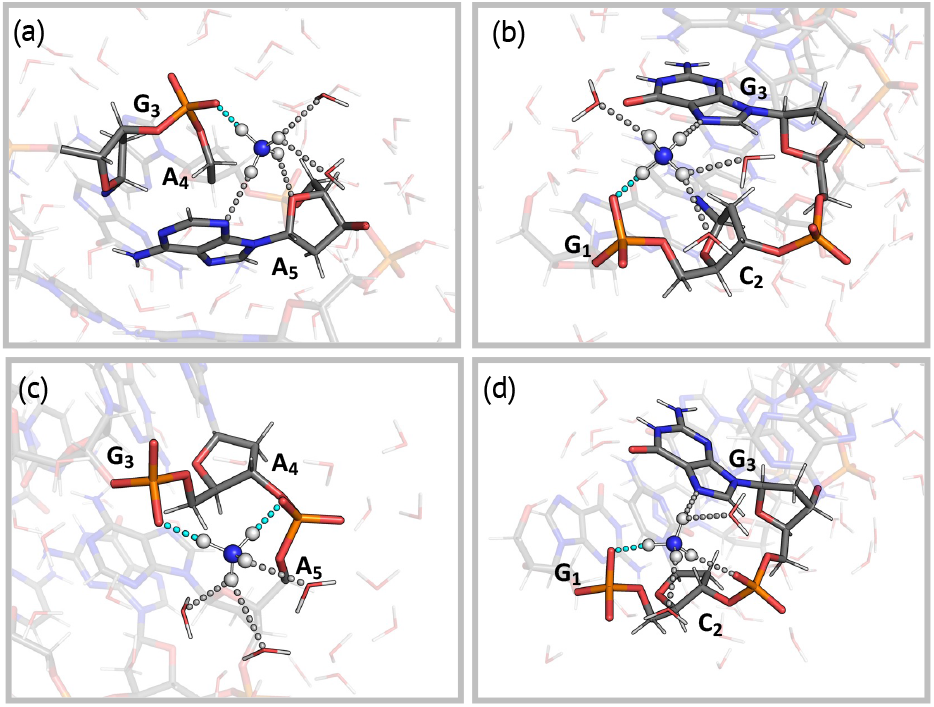
Snapshots of proton transfer configurations showing the interactions of the ammonium ion with (a) the G_3_–p–A_4_ (**IIa-c**), (b) the C_2_–p–G_3_ (**IId-e**), (c) the G_3_–p–A_4_–p–A_5_ (**IIf-h**), and (d) the G_1_–p–C_2_–p–G_3_ (**IIi-j**) moieties. H-bond interactions are shown as dashed lines. The H-bonds involved in the proton transfer are colored in cyan.

### QM/MM MD free energy calculations of complex II

We consider here 10 systems, differing in the hydration level (from ∼80 to ∼30 water molecules, as obtained in our MD simulations, see Tab. 1)^1^ and on whether they interact with one phosphate (interaction mode **(P)**) or with two (interaction mode (**P2**)). We calculate the free energy associated with proton transfer in each mode.

**Table 1:**
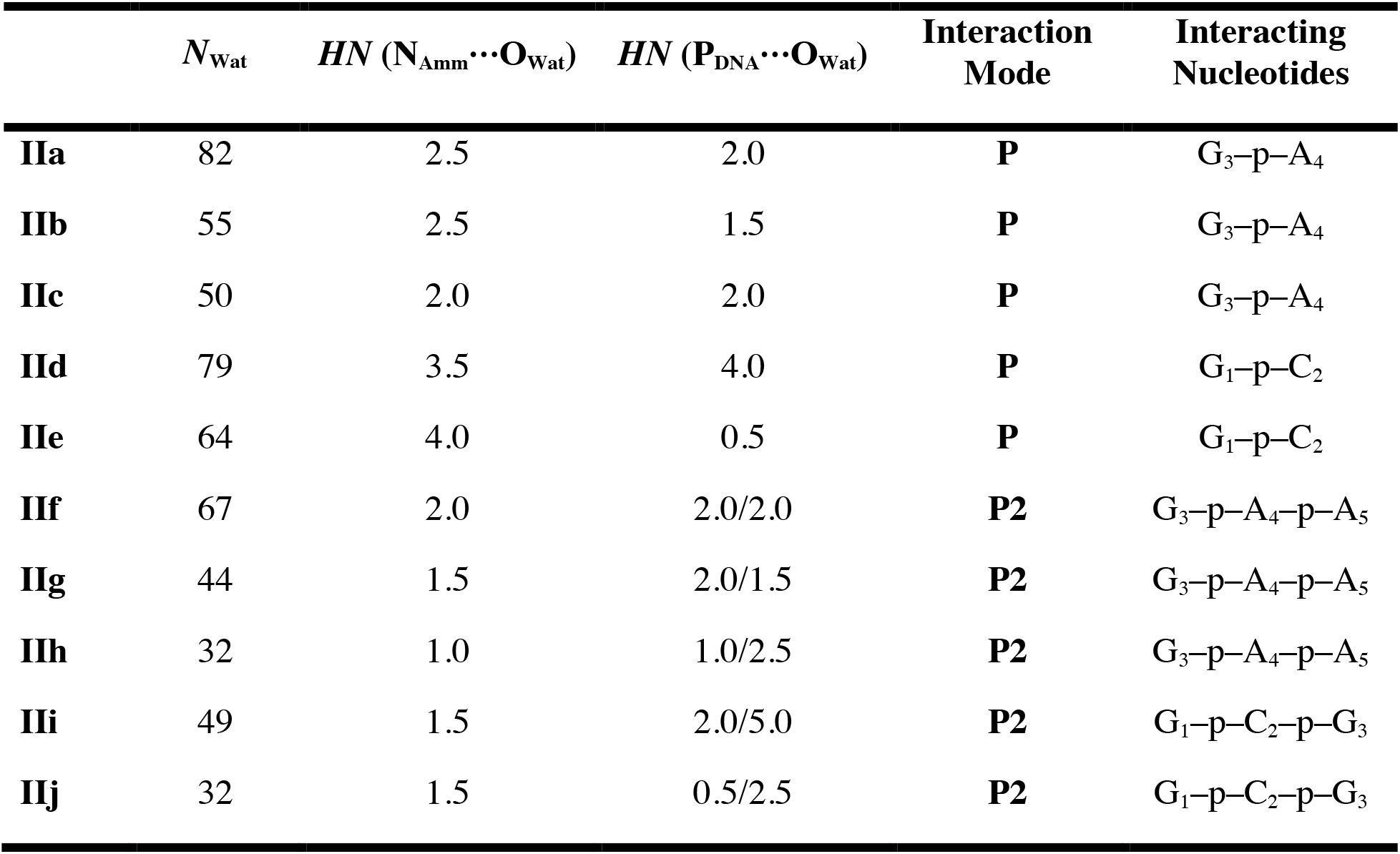
Complex **II** featuring interaction modes **(P)** or (**P2**) at different levels of hydration, undergoing QM/MM simulations. *N*_Wat_ is the total number of water molecules. The *HN* values are calculated from the last 1.25 ps of the equilibration phase. The interaction modes are depicted in Figure 4.

In the first case (Fig. 4a), for *HN* ∼ 2.5, the proton is delocalized between the oligomer and the ammonium (**IIa-b**, Fig. 5 and S16). For *HN* ∼2, the proton is still delocalized between the two moieties, but it mostly sits onto the oligomer (**IIc**, Fig. 5). Thus, decreasing solvation leads the proton towards the oligonucleotide, likely because of the lack of stabilization of the charged ammonium group from the surrounding water molecules.

**Figure 5:**
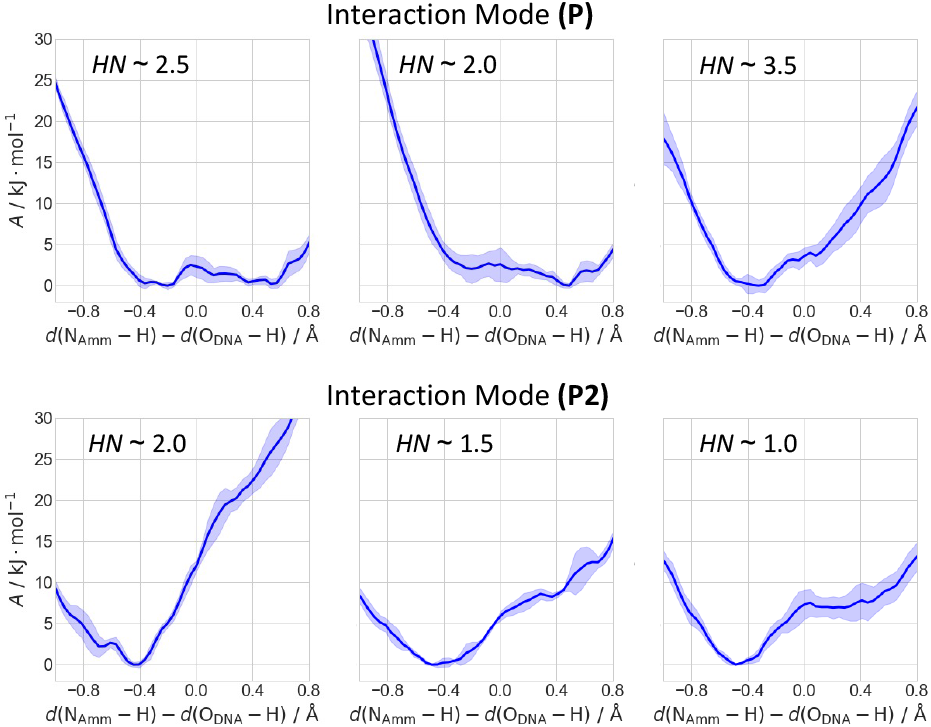
Free energy as a function of the difference of the bond breaking/forming N_Amm_–H and H–O_DNA_ bond distances. Top: Representative profiles of interaction mode (**P**) at different *HNs*, showing systems **IIa** (left), **IIc** (middle) and **IId** (right). Bottom: Representative profiles of interaction mode (**P2**) at different *HNs*, showing the protonation of the G_3_–p–A_4_ moiety in systems **IIf** (left), **IIg** (middle) and **IIh** (right). The complete set of free energy profiles are shown in the SI (Fig. S16).

In the second case (**IId-e** in Fig. 4b), the ammonium cation is markedly more hydrated than in the first (*HN ∼*3.5–4.0, see Tab. 1). As a result, the proton is now *only* localized on the ammonium group (highly stabilized by its solvation shell) and the oligomer is not protonated (Fig. 5).^2^

In the interaction mode **P2**, the ammonium group is less hydrated than in **P**. Either it interacts with the G_3_–p–A_4_–p–A_5_ moiety (**IIf-h**, Fig. 4c) or with G_1_–p–C_2_–p–G_3_ (**IIi-j**, Fig. 4d).^3^ In the first case, it can transfer the proton to either G_3_–p–A_4_ (phosphate “p_3-4_”) or A_4_–p–A_5_ (“p_4-5_”). At *HN* ∼ 2 (the largest value here), the proton is localized on the ammonium group. This contrasts with the interaction mode **P**, where the proton is delocalized. Thus, the interaction of the ammonium group with a second phosphate stabilizes the ionic state. Lowering *HN* slightly increases the probability of p_3-4_ to be protonated (Fig. 5 and Tab. 1; *HN* ∼ 1.5 for **IIg** and ∼ 1 for **IIh**), while the protonation on p_4-5_ remains unlikely (see SI for further details). Thus, the decrease of solvent interactions with the complex only slightly destabilizes the ionic state. In the second case, the proton is localized only at the ammonium group even with *HN* ∼ 1.5. The ionic state is, as described above, stabilized by water and the presence of an additional phosphate.

In conclusions, our simulations suggest that low hydration leads to full protonation of the oligo in interaction mode **P** and it causes an increase of probability to be protonated in **P2**.

In N-ESI-MS experiments of DNA, the evaporation of the droplets leads eventually to protonated oligomers and ammonia.^3,10,14^ The molecular determinants of the process are not known. Here, we address this issue by multiscale simulations on the d(GpCpGpApApGpC) heptanucleotide, for which experimental N-ESI-MS data are available, as well as the model DMP, in the presence of ammonium counterions. The simulations of the model systems (Fig. 2) clearly indicate the impact on hydration on the proton transfer free energy profile: the proton is transferred to DMP if the ammonium is not fully hydrated. MD simulations on the actual heptanucleotide show that the ammonium forms two different interactions with the biomolecule, involving either one (**P** in Fig. 4) or two phosphates (**P2**). Both underwent QM/MM US-based free energy calculations. For **(P)**, where only one phosphate interacts with the ammonium, we find that as soon the ammonium hydration is sufficiently low the proton is transferred to reduce the total charge state of the oligonucleotide. For (**P2**), where two phosphates interact with the ammonium ion, the oligomer is still charged for poorly hydrated ammonium ions, however the probability of being protonated does increase with a decrease of the hydration of the cation. This suggests that the DNA oligomer is eventually protonated when fully dehydrated, consistent with experimental evidence.^14,20^

## COMPUTATIONAL METHODS

### Force fields

The parmBSC1^23^ and TIP3P^24^ force fields were used for the biomolecules (**I** and **II**) and water molecules, respectively. Atomic partial charges for DMP and ammonium ions were derived to be compatible with the employed AMBER force field by a RESP fit at the HF/6-31G* level of theory.^25,26^ Bonded and van-der-Waals parameters for the ammonium ions were taken from the AMBER parm99 library.^27^ The same force field was used for the MM part in the QM/MM calculations.

### MD in aqueous solution

Model **I** consists of DMP and one ammonium ion in a (43.5 × 41.9 × 43.2) Å^3^ water box, containing 5,646 atoms, while **II** consists of the heptamer and six ammonium ions in a (66.7 × 75.9 × 79.1) Å^3^ water box, containing 39,714 atoms. The initial structures were taken from the model with PDB code 1PQT^21^ and solvated with the GROMACS solvate module.^28^ The electrostatic interactions were calculated with the smooth particle mesh Ewald algorithm using a real space cutoff of 10 Å.^29,30^ The same cutoff was employed for the van-der-Waals interactions. The center of mass motion was removed every 100 steps. Bond distances involving covalently bound hydrogen atoms were constrained with the LINCS algorithm.^31^ Periodic boundary conditions and a time step of 1 fs was used throughout. The systems first underwent 5,000 steps of steepest descent minimization. Then, they were heated up to 300 K with 5ns-long simulations at constant volume using the velocity rescaling thermostat algorithm with a coupling constant *τ* of 0.1 ps.^32^ Next, the systems’ density was equilibrated for 5 ns by performing NPT simulations employing the same thermostat algorithm as in the previous step and the Berendsen barostat algorithm (coupling constant of 1 ps) to achieve a pressure of 1 bar.^33^ Finally, production simulations were carried out within the NPT ensemble at 300 K and 1 bar. For this purpose, the Nosé-Hoover thermostat^34,35^ and Parrinello-Rahman barostat^36^ were employed with coupling constants of 0.5 ps and 1.0 ps, respectively. We collected 10 and 500 ns for the (**I**) DMP and (**II**) DNA systems in the water solution, respectively. The following quantities were determined as a function of time: root-mean-square deviation, radius of gyration, and hydrogen bond interactions of the DNA oligo, the distance between NH_4_^+^ ions and phosphate groups for the ion pair characterization and radial distribution functions for the NH_4_^+^ ions and phosphate groups with respect to water oxygen atoms (see SI for details).

### MD and QM/MM in the gas phase

The initial structures were taken from the last snapshots of the MD simulations. For **I** (**II**), we selected three droplets of approximately 800 (3,680) water molecules within a radius of 15 Å (30 Å) around the center of mass of the biomolecules. They also turned out to contain 1 and 3 ammonium ions, and to bear a charge of 0 and –3, respectively for **I** and **II**. Electrostatic interactions were treated by direct Coulomb summation, whereby no cutoff is applied. The same holds for the Lennard-Jones interactions. Both the center of mass translational and rotational velocity was removed every 100 steps. The target temperature was controlled by a velocity rescaling algorithm (*τ* = 0.1 ps).^32^ The systems underwent 50 and 150 ns of NVT simulations at 300 K, following Ref. (19,37), re-assigning the velocities at the beginning of each batch according to the Maxwell-Boltzmann distribution and removing the water molecules located at 60 Å or more from the center of mass of the biomolecules at the end of each simulation. The last sticky water molecules were then evaporated by three consecutive 25ns-long simulations at 350, 400 and 450 K. The same quantities were monitored as above, along with the hydration number (*HN*) of the ammonium and of the phosphate moieties (see SI for details).

The initial QM/MM configurations were taken from the gas phase MD simulations, in which ion pairs between NH_4_^+^ and DNA were observed (see SI for the definition of ion pairs). The systems were inserted in a large box of (200 × 200 × 200) Å^3^. In **I**, the NH_4_^+^ and DMP ions were treated at the QM level, while the water molecules were described at the MM level. In **II**, the ammonium ion and the DNA backbone atoms, which were involved in the studied proton transfer, were enclosed in the QM region (28–41 atoms, Fig. S15). The covalent C4’–C5’ and glycosidic bonds were cut and the dangling bonds saturated with hydrogen link atoms. The QM problem was solved within the density functional theory (see below). Born-Oppenheimer MD and umbrella sampling (US)^38^ was carried out using a time step of 0.5 fs. We used the WHAM analysis to calculate the free energy from the US calculations.^39^

For the simulations of **I**, a planewave basis set up to a cutoff of 100 Ry was used,^40^ with Troullier-Martins pseudopotentials describing the valence shell-core electron interactions.^41^ The PBE exchange-correlation functional was employed.^42^ Periodic images were decoupled from the unit cell with the Martyna-Tuckerman solver.^43^ The system was heated up with a rate of 0.12 K· fs^−1^ to 300 K in 2.42 ps using the Berendsen thermostat (coupling strength of 5,000 a.u.)^33^ and then equilibrated for other 2.42 ps using the Nosé-Hoover thermostat (coupling frequency of 3,500 cm^−1^).^34,35^ The collective variable (CV) for the US calculations was chosen as the difference distance of the breaking/forming N_Amm_–H and H–O_DMP_ bonds (Chart S1). Initial configurations were generated by performing a 2.66ps-long simulation, scanning the CV with a moving restraint. Then, 11 equidistant windows of 14.5 ps in the interval [–1.0; 1.0] Å were simulated. A harmonic restraint with a force constant *k* “ 40 kJ· mol^−1^· Å^−2^ was applied. The first 1.1 ps are discarded from the analysis.

For the simulations of **II**, the quantum problem was solved using the mixed Gaussian-planewave DFT approach.^44,45^ We employed the PBE-D3(BJ)^42,46,47^ functional with the DZVP-MOLOPT basis set^48^ and GTH pseudopotentials.^49^ Four grids were used for the planewave expansion. A density cutoff of 500 Ry was used for the finest grid, and a relative cutoff was set to 80 Ry to specify the coarser grids.^45^ The QM region was electrostatically coupled to the MM potential within the Gaussian expansion of the electrostatic potential approach.^50,51^ After a short minimization, we heated the system to 300 K within 2.5 ps using the velocity rescale algorithm (*τ* = 0.1 ps).^32^ Next, we run an equilibration QM/MM MD at 300 K for 2.5 ps. We finally employed umbrella sampling to predict the free energy for the proton transfer using the same CV as for **I**. 10 equidistant windows of 30.0 ps in the interval [–1.0; 0.8] Å were used (*k* “ 50 kJ· mol^−1^· Å^−2^). The first 5 ps were discarded from the analysis.

The classical MD simulations and the MM simulations in the QM/MM calculations were performed using the GROMACS program package.^28^ The QM calculations were performed either by CP2K^45^ (for **II**, using the API QM/MM interface of GROMACS) or by CPMD^52^ (for **I**, using the MiMiC QM/MM interface).^53–55^ The PLUMED plugin was used to introduce the biases in the umbrella sampling QM/MM simulations.^56,57^

## ASSOCIATED CONTENT

### Supporting Information

Details on the calculated properties; MD analysis of the solution structure of model **II**; additional analysis of the gas phase MD simulations of both models; details on QM/MM-MD re-equilibration; time evolution of all simulated free energy profiles (PDF).

## AUTHOR INFORMATION

### Notes

The authors declare no competing financial interests.

## ACKNOWLEDGMENT

This work was supported by the BioExcel CoE (www.bioexcel.eu), a project funded by the European Union Contract H2020-INFRAEDI-02-2018-823830.

## Footnotes

1 As expected, the larger the number of water molecules, the higher the hydration number values for the ammonium ion and the DNA tend to be.

2 This discussion assumes that the influence of the different nucleobases on the free energy profile is negligible with respect to the influence of the *HN*.

3 **IIi-j** are formed during the QM/MM-MD equilibration phase (see SI).

