## Supporting Information for "Proton Transfers to DNA in Native Electrospray Ionization Mass Spectrometry: A QM/MM Study"

### 1. Calculated Properties and Definitions

Let us consider an atom $i$, either the $NH_4^+$-nitrogen atom ($N_{Amm}$) or the DMP/ heptanucleotide phosphorus atom ($P_{DMP}$ /$P_{DNA}$) coordinated by a group of atoms $j$ (water oxygens ($O_{Wat}$)). Then, the hydration number for atom ($i$) reads:

$$HN(i) = \sum_j \frac{1-\left(\frac{r_{ij}-d_0}{r_0}\right)^6}{1-\left(\frac{r_{ij}-d_0}{r_0}\right)^{12}} \qquad (1)$$

where $r_{ij}$ corresponds to the distance between atoms $i$ and $j$. The parameters $d_0$ and $r_0$ are chosen in the way that only the first peak of the $N_{Amm}$–$O_{Wat}$ or the $P_{DMP/DNA}$–$O_{Wat}$ radial distribution functions is fully included. If $i = N_{Amm}$, $d_0 = 3.0$ Å and $r_0 = 1.0$ Å (Fig. S1a), if $i = P_{DMP}$, $d_0 = 3.7$ Å and $r_0 = 1.0$ Å (Fig. S1b). If $i = P_{DNA}$, $d_0$ and $r_0$ have identical values (see Fig. S1c).

We employed the GROMACS `hbond` module to determine the number of internal hydrogen bonds of the heptanucleotide.[1,2] Cutoffs of 3 Å and 150° was used for the hydrogen-acceptor distance and the donor-hydrogen-acceptor angle.

We defined the ammonium and the anions if the distance between $N_{Amm}$ and $P_{DMP}$ is 4 Å or lower (in the case of DMP) or the distance between $N_{Amm}$ and any $P_{DNA}$ is 4 Å or lower (in the case of the heptamer).

The definition of the collective variable (CV) for all umbrella sampling (US) performed here is shown in Chart S1.

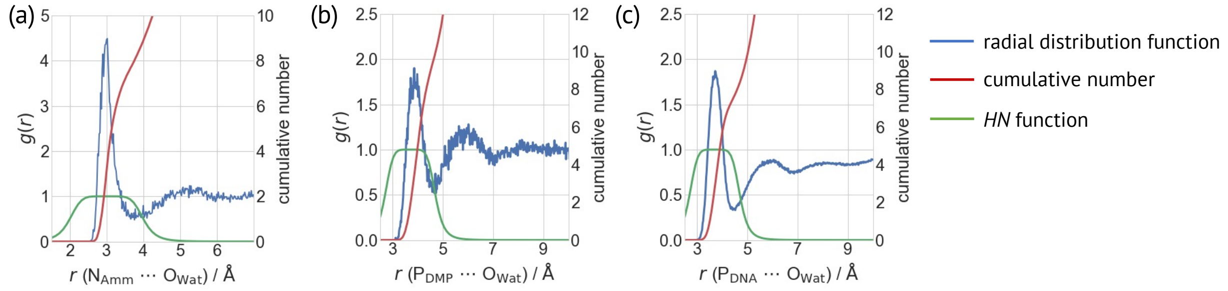


**Figure S1:** Radial distribution functions *g(r)*, cumulative number and summand of equation (1). (a) $N_{Amm}$–$O_{Wat}$ $g(r)$. (b) $P_{DMP}$–$O_{Wat}$ $g(r)$. (c) $P_{DMP}$–$O_{Wat}$ $g(r)$.

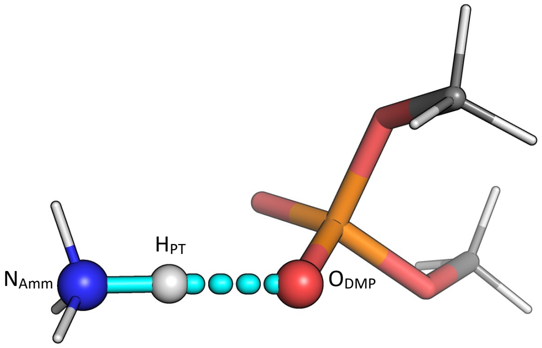


$$\mathrm{CV} := \Delta d = d(\mathrm{N_{Amm}} - \mathrm{H_{PT}}) - d(\mathrm{O_{DMP}} - \mathrm{H_{PT}})$$

**Chart S1.**

### 2. Results: Additional Details

#### 2.1 MD of I in aqueous solution

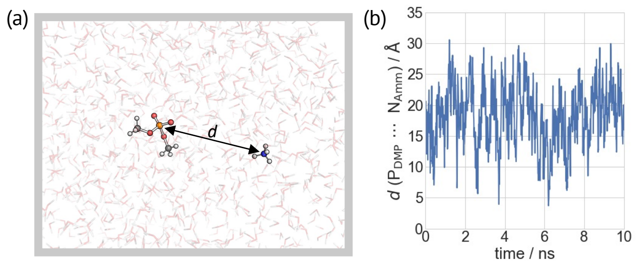


**Figure S2:** (a) Snapshot of **I** in solution, highlighting the $P_{DMP} - N_{Amm}$ distance $d$. (b) $d$ plotted as a function of simulated time.

### 2.2 MD of I in aqueous droplets

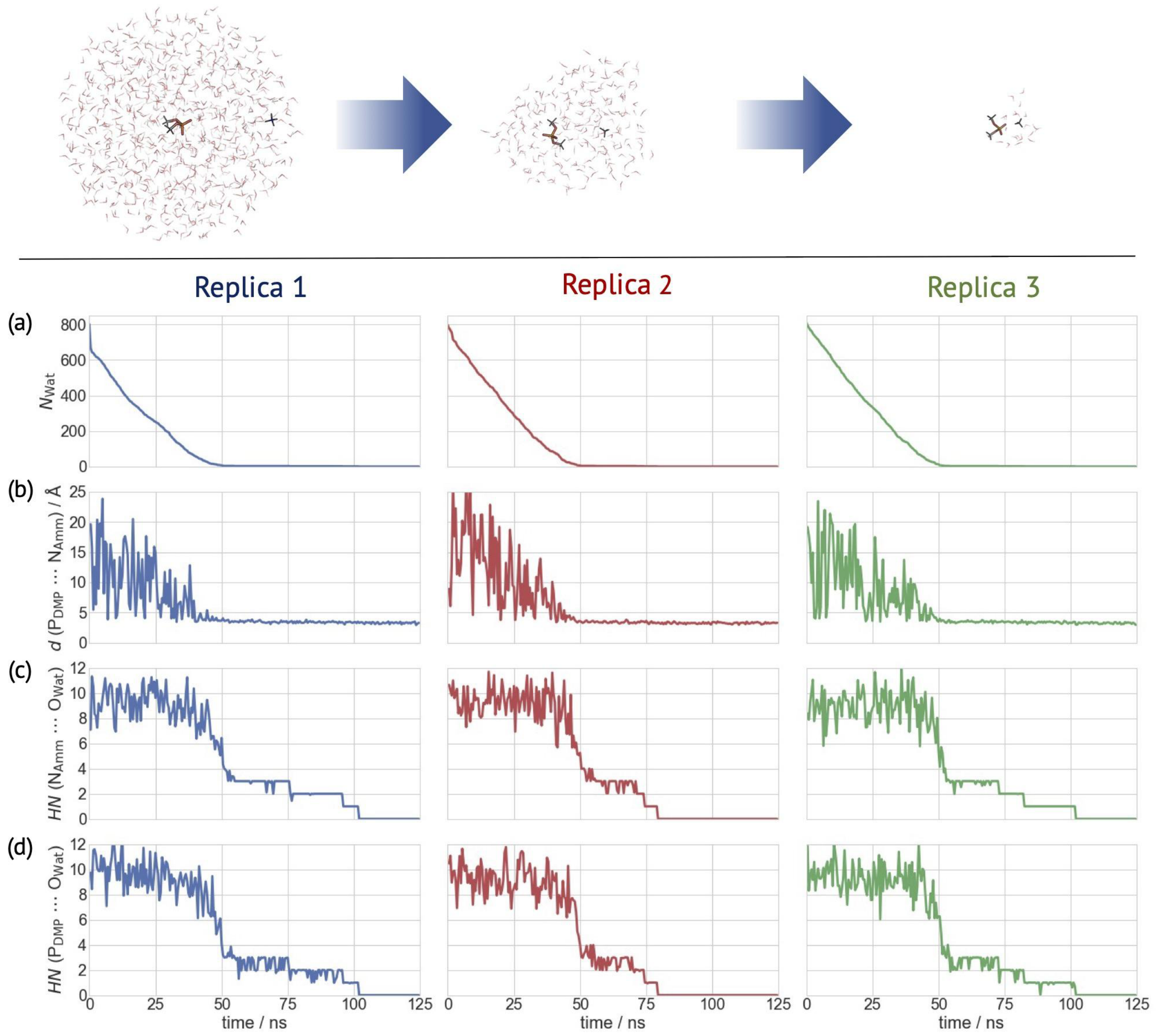


**Figure S3: Top**: Evaporation process of the dimethyl phosphate ion and an ammonium ion in a water droplet as observed during one of our three replicas of the simulation. **Bottom**: Selected properties of the three replicas of gas phase MD simulations plotted as a function of time: **(a)** Number of water molecules in the system ($N_{Wat}$). **(b)** $P_{DMP}$ -$N_{Amm}$ distance; **(c)–(d)** *HN(i)* ($i$ = $N_{Amm}$ **(c)** and $P_{DMP}$ **(d)**).

### 2.3 QM/MM of I in the Gas Phase

We report here the results from 2.42 ps QM/MM MD simulations of **Ia-c.**

The simulation of **Ia** leads readily to proton transfer. The resulting neutral state is stable with *HN* ~ 2-3 for the rest of the dynamics ($d(N_{Amm}–H_{PT})$ = 1.4 ± 0.2 Å, $d(O_{DMP}–H_{PT})$ = 1.2 ± 0.1 Å).

In the QM/MM MD of the other two complexes, the ammonium ion is more hydrated (*HN*s ~ 3–4 and 4–6). No proton transfer is observed. The N–H bond forming an H-bond with DMP ion increases by ~0.1 Å relative to the others (see Fig. S4).

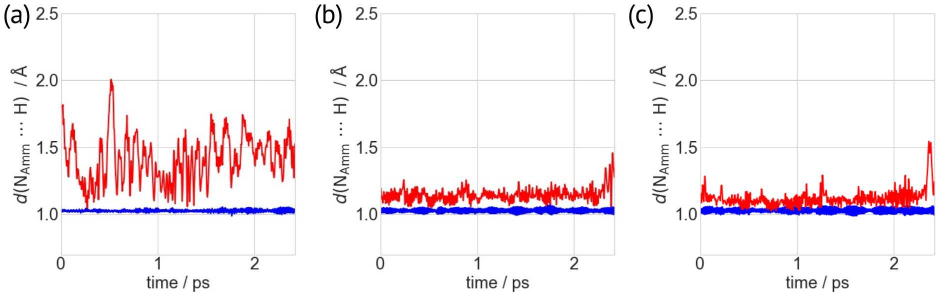


**Figure S4:** $N_{Amm}$–H bond distances of gas phase QM/MM-MD simulations plotted as a function of time for systems (a) **Ia**, (b) **Ib** and (c) **Ic**. The N–H bond involved in the proton transfer is shown in red, while the other N–H bonds are shown in blue.

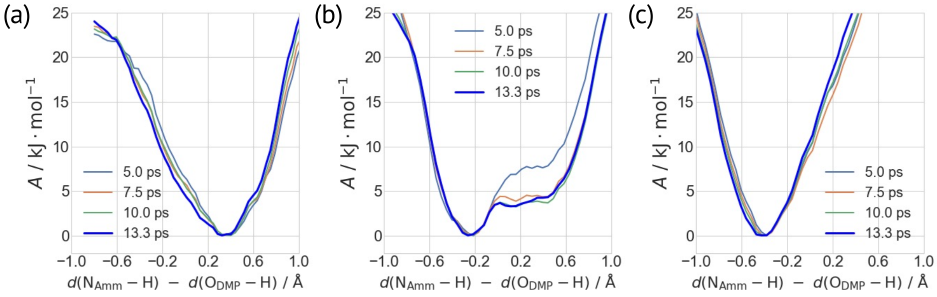


**Figure S5:** Time evolution of the free energy profiles as a function of the difference of the breaking/forming $N_{Amm}$–H and H–$O_{DMP}$ bond distances for systems (a) **Ia**, (b) **Ib** and (c) **Ic**.

### 2.4 MD of II in water solution

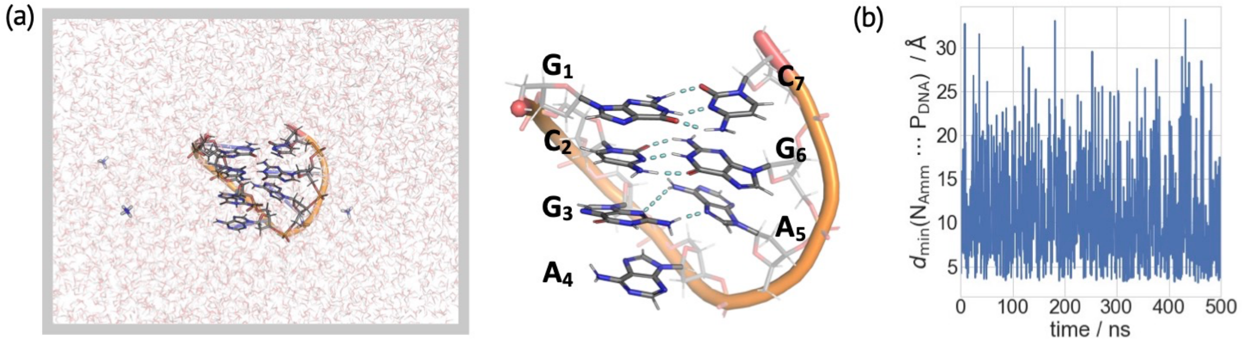


**Figure S6** (a) MD snapshot of complex **II** in aqueous solution, showing the hairpin structure of the heptanucleotide and 6 $NH_4^+$ ions. Inset: zoom on the structure of the heptanucleotide. The H-bonds between base pairs are shown as cyan, dashed lines. (b) $P_{DNA}$–$N_{Amm}$ distance *d* plotted as a function of time.

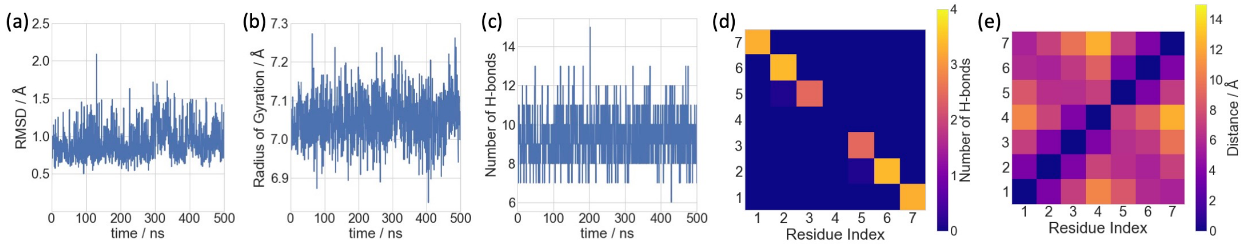


**Figure S7: (a)–(c)** Selected structural descriptors of the heptanucleotide plotted as a function of simulated time. **(a)** Root-mean-square-deviation (RMSD) of all atoms relative to the initial structure. **(b)** Radius of gyration. **(c)** Total number of internal H-bonds. (**d)** Heatmap of the average number of observed hydrogen bonds between the base pairs, reflecting the formation of two canonical base pairs ($G_1$–$C_7$ and $C_2$–$G_6$ with ~3 H-bonds) and one non-canonical base pair ($G_3$–$A_5$ with ~2 H-bonds). **(e)** Contact map of the nucleobases, showing the characteristic X-shape of an antiparallel duplex.[3]

### 2.5 MD of II in the water droplets

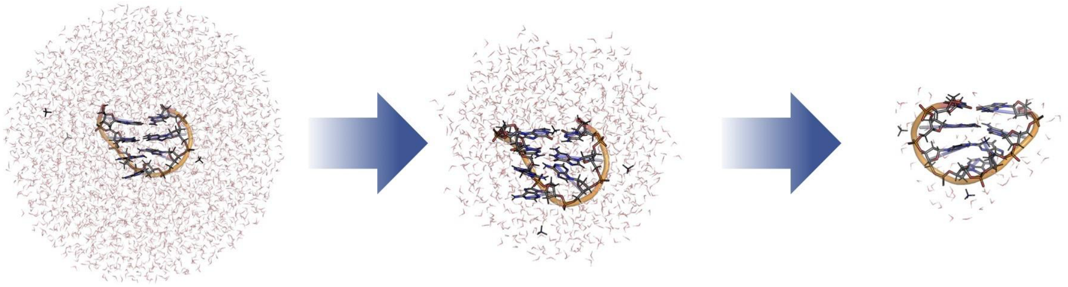

**Figure S8:** Evaporation process of the heptanucleotide and ammonium ions in a water droplet as observed in one of the 225 ns MD simulations performed here.

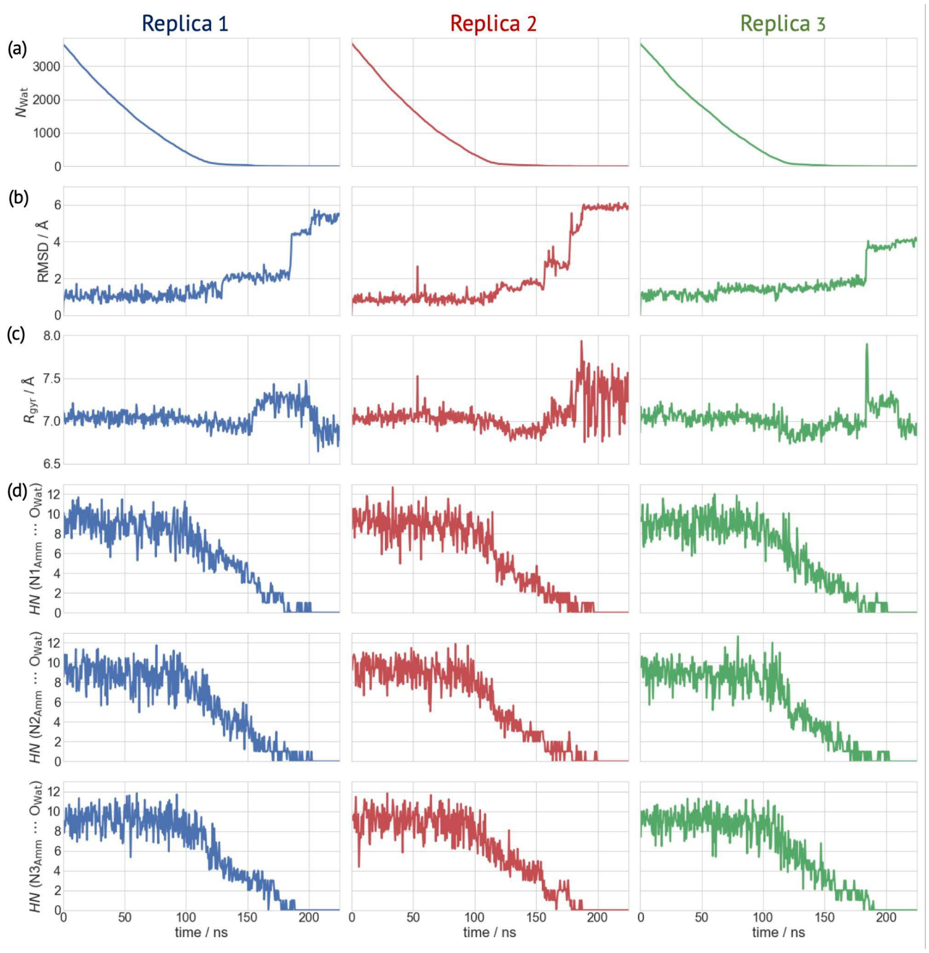


**Figure S9:** Selected properties plotted as a function of simulated time in the three replicas of gas phase MD simulations. (a) Number of water molecules ($N_{\mathrm{Wat}}$). (b) RMSD of heptanucleotide's all atoms relative to the initial structure. (c) $R_{\mathrm{gyr}}$. (d) *HN* of the three ammonium ions.

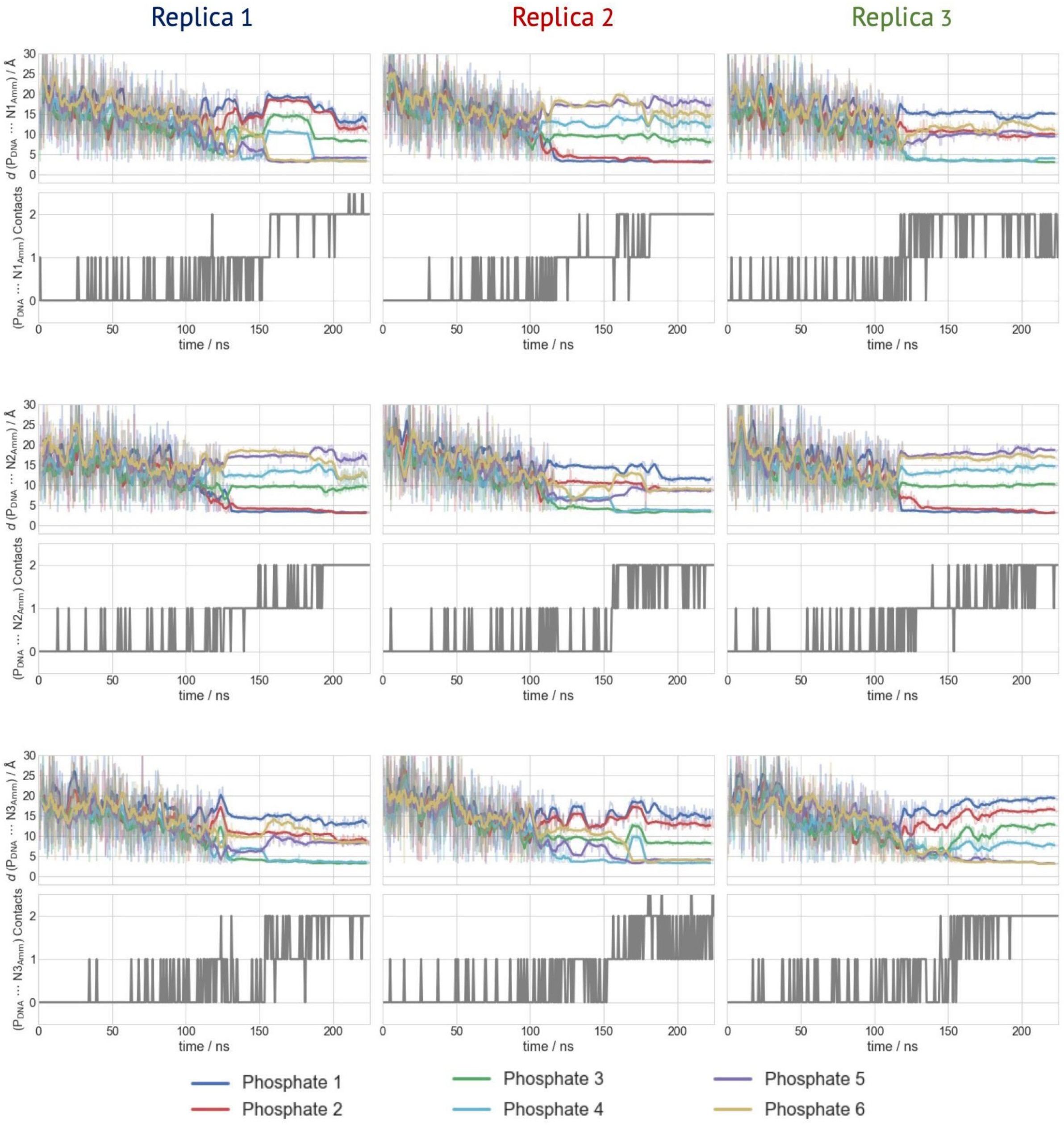


**Figure S10:** Distances *d* between $P_{DNA}$ and the three $N_{Amm}$-atoms from the three MD replicas. Data at the end of each 500 ps batch of MD simulations are shown as transparent lines, while their moving averages (window size of 10) are represented as bold lines. The corresponding number of contacts are shown below each distance plot.

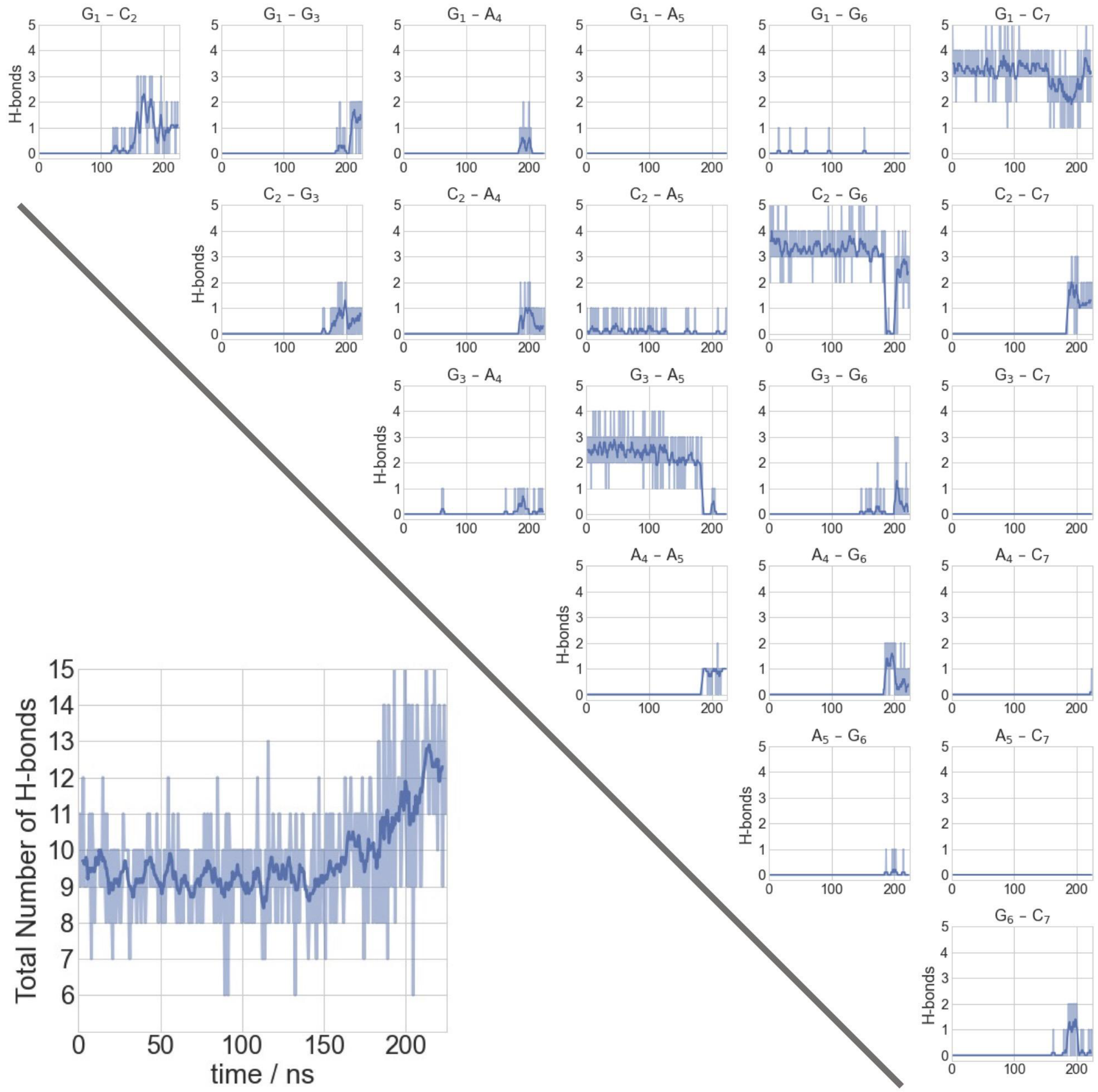


**Figure S11:** Nucleobase H-bond interactions plotted as a function of time (replica 1). Data at the end of each 500 ps batch of MD simulations are shown as transparent lines, while their moving averages (window size of 10) are represented as bold lines. Upper right: Base pair specific number of H-bonds. Lower left: Total number of H-bonds of the heptanucleotide.

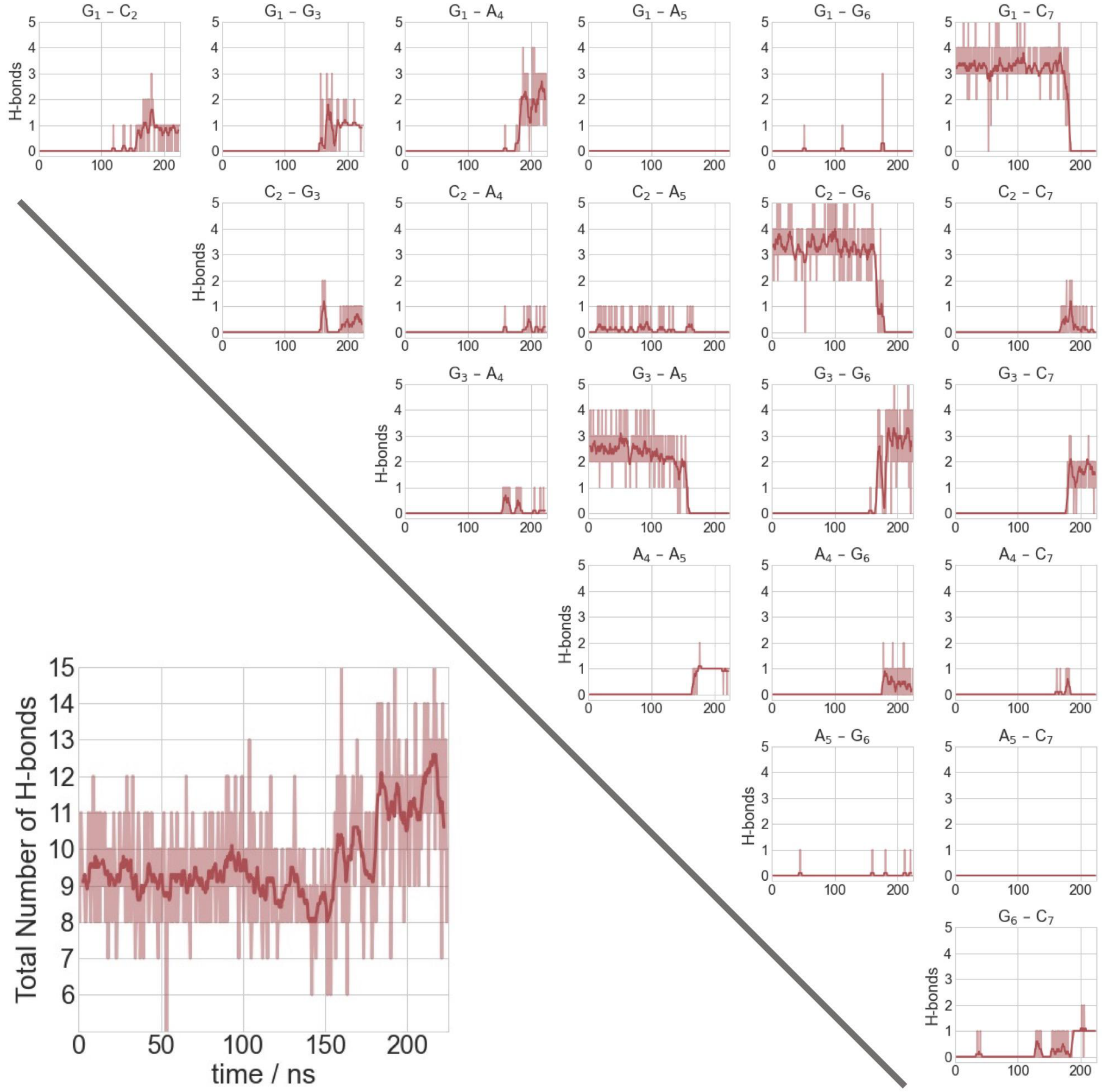


**Figure S12:** Same as Fig. S11 for the second replica of the MD.

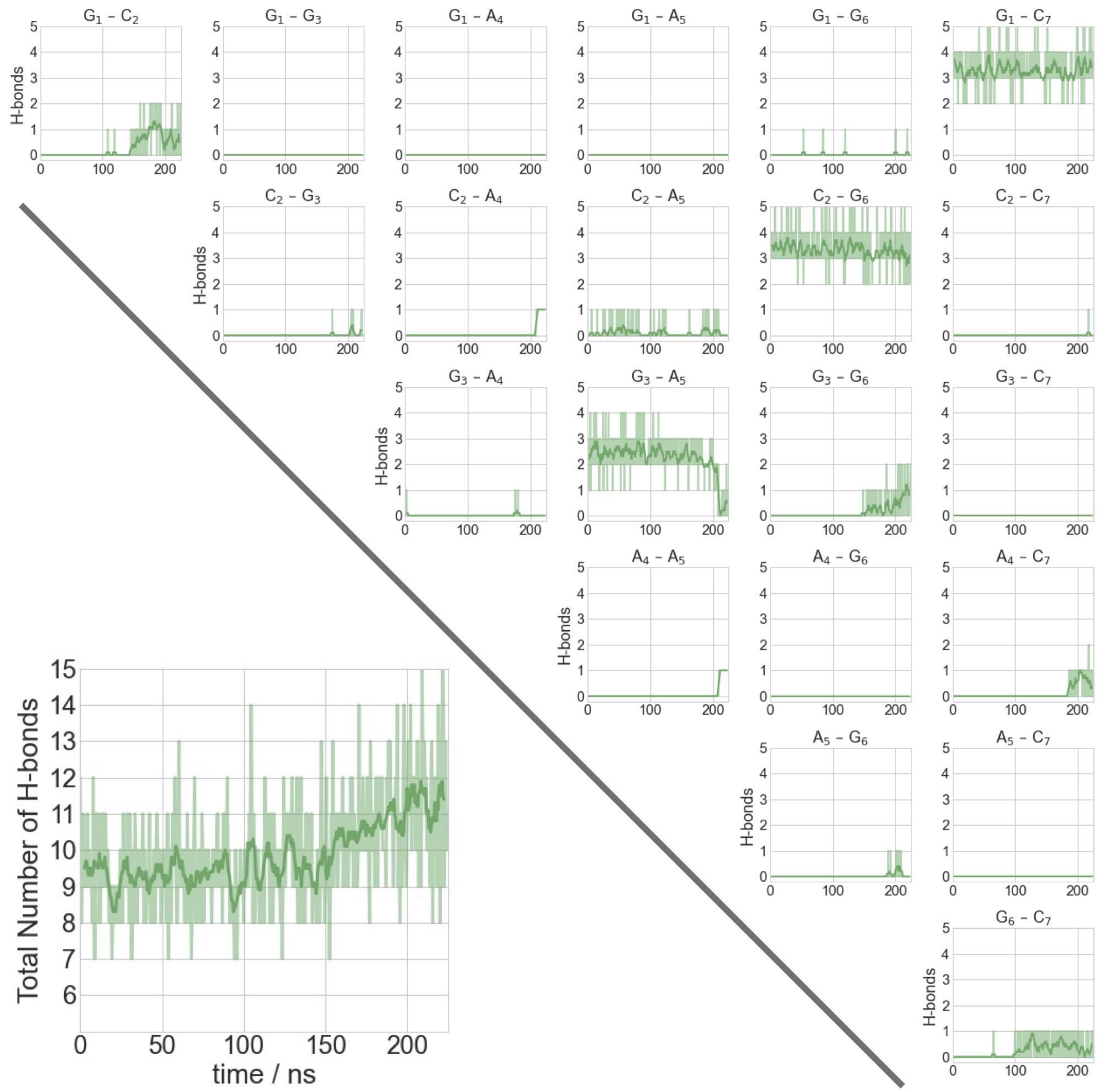


**Figure S13:** Same as Fig. S11 for the third replica of the MD.

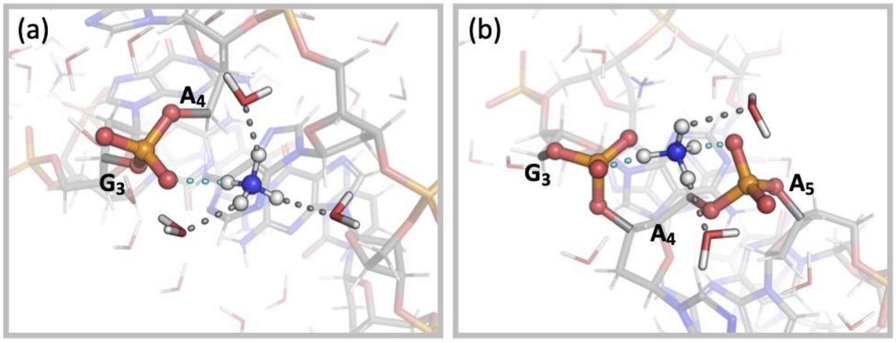


**Figure S14:** Salt bridges between a $NH_4^+$ ion with (a) one phosphate or (b) two phosphate groups as observed in our MD simulations.

### 2.6 QM/MM of II in the Dehydrated Water Droplet

#### 2.6.1 Partitioning of the QM Region for the QM/MM Calculations.

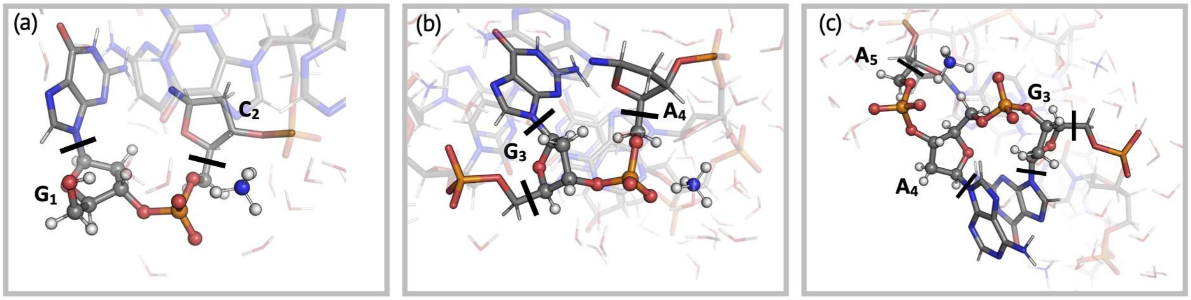


**Figure S15:** QM atoms (spheres) in the QM/MM calculations. Cut covalent bonds indicated as black lines.

#### 2.6.2 QM/MM Simulations of Complexes IIa-j.

We performed 2.5 ps QM/MM MD and 25.0 ps QM/MM US calculations for each complex.

In the 2.5ps-long QM/MM MD of **IIa, IIb** and **IIc,** the ammonium ion forms a H-bond to the phosphodiester group between $G_3$ and $A_4$ (Fig. 4a), with different initial conditions (see Table 1). The $NH_4^+$ ion forms H-bonds to the O4'-deoxyribose and N3- adenine atoms of the $A_5$ residue of the heptanucleotide (Fig. 4a). In **IIa** and **IIb,** no proton transfer is observed. The N–H bond involved in the proton transfer is elongated to 1.2 ± 0.1 Å (**IIa**) and 1.12 ± 0.04 Å (**IIb**) relative to the other bonds of the $NH_4^+$ ion (~ 1.05 Å), while the O–H distance is 1.4 ± 0.1 Å (**IIa)** and 1.5 ± 0.1 Å (**IIb**). The ammonium *HN* fluctuates around ~2–3 (**IIa)** and ~2–4 (**IIb**). In **IIc**, the proton is almost shared between the two moieties. The N–H and O–H distances are observed to strongly fluctuate with 1.3 ± 0.2 Å and 1.4 ± 0.2 Å. The $NH_4^+$ *HN* fluctuates around ~2.

In **IId** and **IIe,** the ammonium ion forms a H-bond to the phosphodiester group between $G_1$ and $C_2$, with different initial conditions (Fig. 4b). No proton transfer is observed. The N–H bond is 1.12 ± 0.05 Å (**IId**) and 1.2 ± 0.2 Å (**IIe),** while the O–H distance is 1.5 ± 0.1 Å (**IId)** and 1.2 ± 0.1 Å (**IIe**). The $NH_4^+$ *HN* strongly fluctuates from ~3–6 (**IId)** and ~2–5 (**IIe**). The $NH_4^+$ ion also forms a H-bond with N7-guanine atoms of the $G_3$ residue of the heptanucleotide (Fig. 4b).

In **IIf**, the ammonium ion forms H-bonds to two phosphodiester groups of the moiety $G_3$–p–$A_4$–p–$A_5$ (Fig. 4c). No proton transfer is observed. The N–H bonds involved in H-bond interactions are just slightly elongated to about 1.09 ± 0.04 Å, while the O–H distances are 1.6 ± 0.1 Å. The $NH_4^+$ *HN* fluctuates around ~2.

In **IIg**, the same interaction mode as for **IIf** is studied, but with different initial conditions (see Table 1). No proton transfer is observed. The ammonium interacts stronger with the $G_3$–p–$A_4$ group than with the $A_4$–p–$A_5$ group: The N–H bonds involved in H-bond with the phosphate groups are 1.2 ± 0.1 Å and 1.1 ± 0.1 Å, respectively. The corresponding O–H distances are 1.5 ± 0.2 Å and 1.7 ± 0.3 Å, respectively. The $NH_4^+$ *HN* fluctuates around ~1–2. The $NH_4^+$ ion features no other H-bond interactions with the heptanucleotide as in **IIf**.

In **IIh** exhibits the same interaction mode as for **IIf** and **IIg** but with different initial conditions (see Table 1). No proton transfer is observed. However, the ammonium interacts strongly with the $G_3$–p–$A_4$ group: The N–H and O–H bond lengths are very similar with 1.3 ± 0.2 Å and 1.3 ± 0.2 Å, respectively. The interaction to the $A_4$–p–$A_5$ group leads to a slightly elongated N–H bond with

1.07 ± 0.04 Å, while the O–H distance is 1.7 ± 0.2 Å. The $NH_4^+$ *HN* fluctuates around ~0–2. The $NH_4^+$ ion features no other H-bond interactions with the heptanucleotide.

**IIi** shows the same interaction of the $NH_4^+$ ion with the G1–p–C2 group as for **IId** and **IIe,** but with different initial conditions (see Table 1). During the dynamics, the ammonium ion additionally forms a H-bond to the phosphodiester group between $C_2$ and $G_3$ (Fig. 4d). No proton transfer is observed. The N–H bond involved in proton transfer is 1.1 ± 0.1 Å, while the O–H distance is 1.5 ± 0.1 Å. The $NH_4^+$ *HN* varies from ~2–5.

In **IIj**, the same interaction mode evolves as for **IIi** (Fig. 4d), but from different initial conditions (see Table 1). The N–H bond involved in proton transfer is elongated to 1.2 ± 0.2 Å relative to the other bonds of the $NH_4^+$ ion (~ 1.05 Å), while the O–H distance is 1.3 ± 0.2 Å. The $NH_4^+$ *HN* varies from ~1–2.

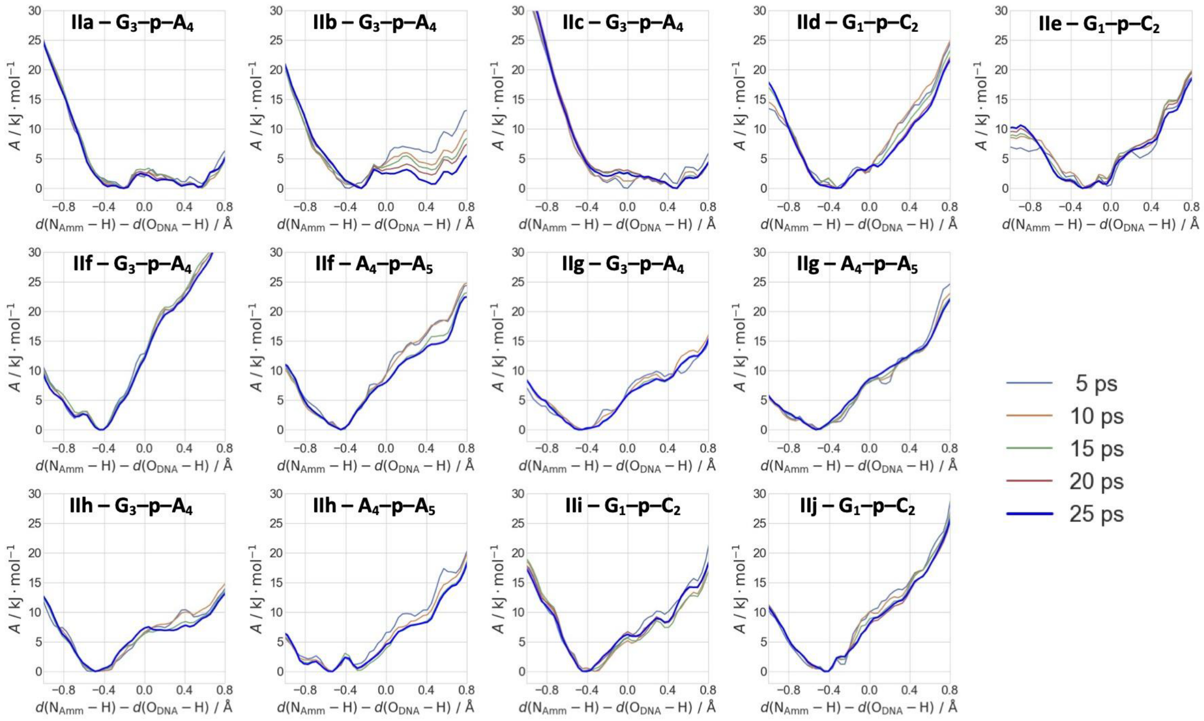


**Figure S16:** Time evolution of the free energy profiles as a function of the difference of the breaking/forming $N_{Amm}$–H and H–$O_{DNA}$ bond distances. Details on the systems and the interaction modes are described in the main text (see Fig. 4 and Table 1).
